# Studying the drug treatment pattern based on the action of drug and multi-layer network model

**DOI:** 10.1101/780858

**Authors:** Liang Yu, Yayong Shi, Quan Zou, Lin Gao

**Affiliations:** School of Computer Science and Technology, Xidian University, Xi’an, 710071, P.R. China; Institute of Fundamental and Frontier Sciences, University of Electronic Science and Technology, 650004, P.R. China

**Keywords:** Drug Treatment Pattern, Drug-target Modules, Multi-layer Network, Tissue Specificity, Action of Drug

## Abstract

**Objectives:** A drug can treat multiple diseases, indicating that the treatment of the drug has certain patterns. In this paper, we studied the treatment pattern of drugs from a new perspective based on the ***a***ction of drug and ***m***ulti-layer network model (STAM). Diseases affect the gene expression in related tissues and each disease corresponds to a tissue-specific protein-protein interaction (TSPPI) network. Therefore, a drug is associated with a multi-layer TSPPI network associated with diseases it treats. Single tissue-specific PPI network cannot consider all disease-related information, leading to find the potential treatment pattern of drugs difficultly. Research on multi-layer networks can effectively solve this disadvantage. Furthermore, proteins usually interact with other proteins in PPI to achieve specific functions, such as causing disease. Hence, studying the drug treatment patterns is equivalent to study common module structures in the multi-layer TSPPI network corresponding to drug-related diseases. Knowing the treatment patterns of the drug can help to understand the action mechanisms of the drug and to identify new indications of the drug.

**Methods:** In this paper, we were based on the action of drug and multi-layer network model to study the treatment patterns of drugs. We named our method as STAM. As a case of our proposed method STAM, we focused on a study to trichostatin A (TSA) and three diseases it treats: leukemia, breast cancer, and prostate cancer. Based on the therapeutic effects of TSA on various diseases, we constructed a tissue-specific protein-protein interaction (TSPPI) network and applied a multi-layer network module mining algorithm to obtain candidate drug-target modules. Then, using the genes affected by TSA and related to the three diseases, we employed Gene Ontology (GO), the modules’ significance, co-expression network and literatures to filter and analyze the identified drug-target modules. Finally, two modules (named as M17 and M18) were preserved as the potential treatment patterns of TSA.

**Results:** The processed results based on the above framework STAM demonstrated that M17 and M18 had strong potential to be the treatment patterns of TSA. Through the analysis of the significance, composition and functions of the selected drug-target modules, we validated the feasibility and rationality of our proposed method STAM for identifying the drug treatment pattern.

**Conclusion:** This paper studied the drug treatment pattern from a new perspective. The new method STAM used a multi-layer network model, which overcame the shortcomings of the single-layer network, and combined the action of drug. Research on drug treatment model provides new research ideas for disease treatment.

## 1 Introduction

Drugs interact with the target and off-target, thus triggering the downstream signal cascade, leading to disturbance in the cell’s transcriptome [1]. Discovering new drug targets is critical and helpful for deep understanding the drug mechanistic action [2], which is essential to drug discovery [3–6], clinical trials, and efforts to overcome drug resistance [7, 8]. Researchers have devoted themselves to the study of specific mechanisms between drug and targets [9–15]. Drug target is not limited to single gene, but also include modules or pathways that participate in the regulation of disease processes, named as drug-target modules [16]. Although each gene in a drug-target module may not contribute to the treatment of the disease, when they act together, they play a significant role in the treatment of the disease [16]. By targeting genes in a drug-target module, it becomes possible to identify the function at the level of biological tissue associated with complex disease pathologies [17, 18].

One drug, one disease. This is how we traditionally consider drugs, but many of them are actually effective against many diseases [19], which means that the drug treatment has a certain pattern. In this paper, we attempted to study drug treatment patterns through drug-target modules in a multi-layer network. Genes in drug-target modules are highly expressed in disease-associated tissues [20–22], which allows drug targets to be treated by specific small molecular compounds [23–26]. Each disease corresponds to a tissue-specific protein-protein interaction (TSPPI) network [27]. Therefore, a drug may be associated with a multi-layer TSPPI network. Research exploring bio-molecular networks has gradually shifted focus from single-layer networks to multi-layer networks [28] because biological networks complement each other [29, 30]. This allows a deep understanding of the treatment patterns of the drug by extracting target modules from the multi-layer TSPPI network associated with the drug.

We proposed a new perspectives for studying drug treatment patterns based on the action of drug and multilayer network model (STAM). Figure 1 provides the framework of our proposed method, STAM. Here, we took drug trichostatin A (TSA) [31] and three diseases it treated, leukemia, breast cancer and prostate cancer [32–36], as a case for studying. Finally, we identified two drug-target modules of TSA (M17, M18) as the potential treatment patterns of TSA.

**Figure 1:**
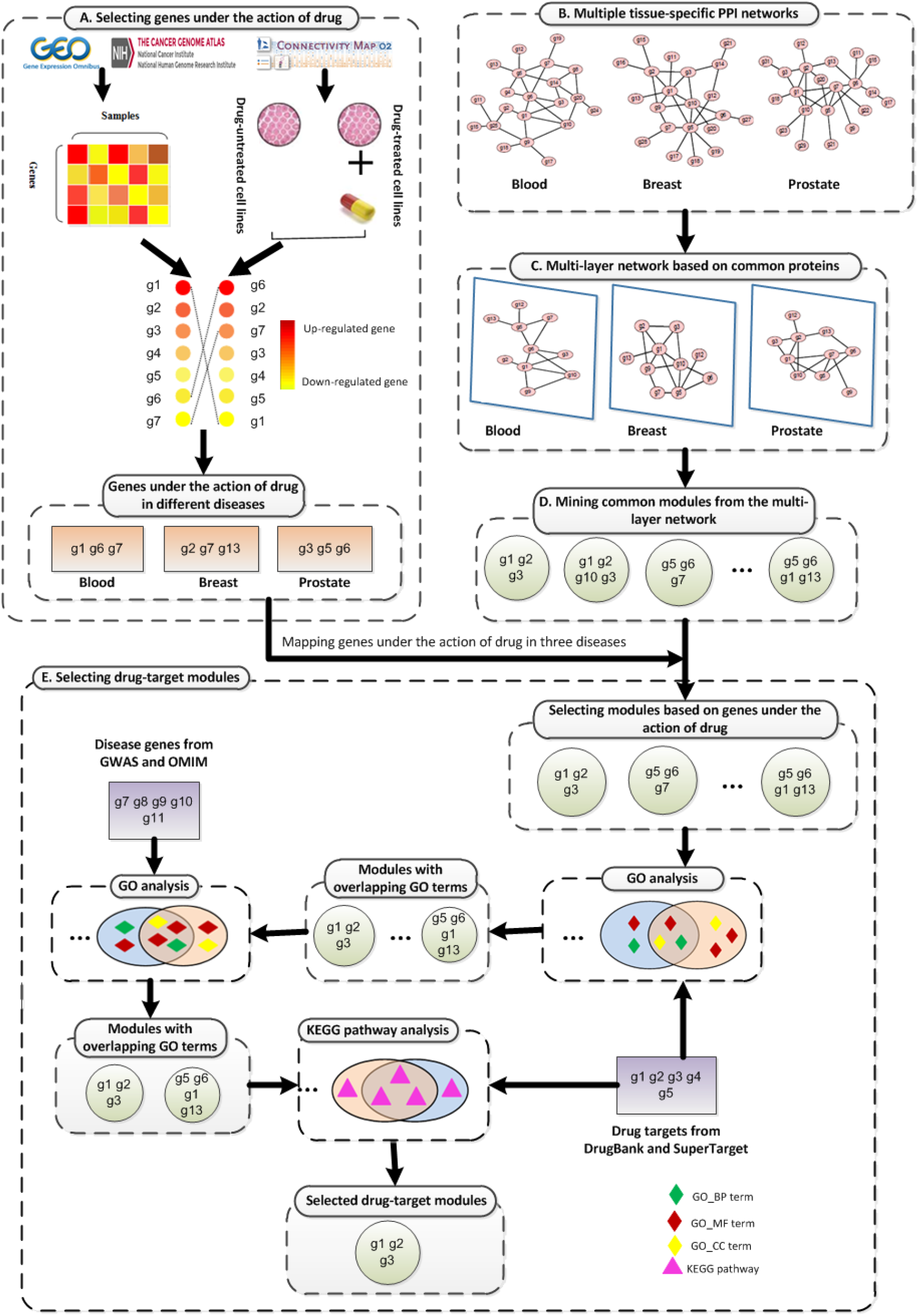
The framework of our method STAM to study the drug treatment pattern through extracting drug-target modules from a multi-layer tissue-specific protein-protein interaction (TSPPI) network. A. Selecting genes under the action of drug trichostatin A (TSA) on three diseases it treated based on gene differential expression profiles. B. Processing TSPPI networks related to three diseases from GIANT database. C. Standardizing a multi-layer spanning subgraph based on common proteins got from step B. D. Using tensor-based mining algorithm to identify drug-target modules from the multi-layer TSPPI network. E. Selecting drug-target modules and validating them. Finally, the preserved modules were likely to be the treatment patterns of the drug.

The remainder of this paper is organized as follows. Section 2 presents an overview of related work. Section 3 presents our data and a description of the proposed method STAM. Section 4 provides details of our empirical work and our results. The conclusion is given in Section 5.

## 2 Related Work

Over time, researchers have worked toward maintaining an accurate and up-to-date map of approved drugs and their targets. In 1996, Drews and Ryser estimated the number of human molecular targets for approved small-molecule drugs [37]. Then Hopkins and Groom [38] were the first to attempt to define the druggable genome based on existing successful drug development programs. Subsequent analyses [39–44] led to the concept of “privileged” protein families, an idea that has contributed to a consistent and successful history of drug discovery.

The original method for the study of drug targets was based on ligands because ligands with chemical similarity are involved in similar chemical activities and can bind similar targets [45]. Compared to the traditional methods of accidental discovery, approaches based on targets can find more valuable drugs. This kind of approach is aimed at stimulating the affinity between ligands and protein targets. Wang et al. [9] designed a multi-target drug complex to disrupt the common network structure of four cancers using a virtual screening approach based on ligands and structure.

Advances in high-throughput protein sequencing technology brought increasing improvements to protein interactions, making identification of drug targets more accurate. For example, Vilar et al. [10] found a new molecular drug by calculating the similarity of protein interaction spectrums. McCormick [11] studied the interactions of K-Ras through post-translational modification and signaling proteins, providing more treatment opportunities for cancers. Kaltdorf et al. [12] established a metabolic network model using basic pattern analysis and flow approximation. They analyzed the specific enzymes of pathogens, and finally obtained multiple antifungal target sites.

Wang et al. [46] applied the seed-connector algorithm to find a placebo treatment response module in the PPI network. By calculating the shortest distance, this module was found to be close to the disease module and drug target. Through a series of molecular pathway analyses, this module could be used as a target module for drugs. This study provided a better understanding of the molecular mechanisms of the placebo response, set the stage for minimizing its impact in clinical trials, and promoted strategies for its use in treatment.

## 3 Data and Method

To develop and validate our proposed method STAM for studying drug treatment pattern based on the action of drug and multi-layer network model, we took trichostatin A (TSA) and three diseases it treated (leukemia, breast cancer and prostate cancer) as a case.

### 3.1 Datasets

#### Gene expression data under the action of TSA

From the Connectivity Map (CMAP) [47, 48], we downloaded the gene expression data of multiple samples under the action of TSA. These data were measured in the MCF7, PC3, and HL60 cell lines using HT_HG-U133A chips. Meanwhile, we downloaded the raw data of the 83 MCF7 cell lines, 52 PC3 cell lines, and 25 HL60 cell lines.

#### Gene expression data under disease state

We downloaded the gene expression profiles of breast cancer and prostate cancer from The Cancer Genome Atlas (TCGA) [49]. The dataset of breast cancer contains 1212 samples, including 1100 tumor samples and 112 control samples. The dataset of prostate cancer contains 550 samples, including 498 tumor samples and 52 control samples. We downloaded the gene expression profiles of leukemia from the NCBI Gene Expression Omnibus (GEO) [50]. The gene expression dataset (GSE48558) contains 170 samples, including 121 tumor samples and 49 control samples.

#### Tissue-specific PPI networks

We downloaded tissue-specific PPI networks marked as “top edges” from the Genome-scale Integrated Analysis of Gene Networks in Tissue (GIANT) database [27]. “Top edges” signifies that the network is filtered to include only the edges that have evidence supporting a tissue-specific functional interaction. The GIANT database integrated diverse genome-scale data in a tissue-specific manner to construct 144 human tissue- and cell lineage-specific networks. This work demonstrated their broad usefulness for generating specific testable hypotheses, summarizing tissue-specific relationships between diseases, and reprioritizing results from genetic association studies. In the meantime, we downloaded tissue-specific PPI networks of breast, prostate, and blood tissues.

### 3.2 Preprocessing gene expression data

The gene expression data was downloaded from CMAP. The filename suffix is .CEL. The .CEL files record the signal intensity of each probe. In this paper, we used the RMA package of R software to standardize and visualize the data from the .CEL files. We mapped probes to genes based on the existing available platform information. If a probe failed to map to any gene, the probe and its corresponding expression data were deleted. If multiple probes were mapped to a single gene, the expression values of multiple probes were averaged as the expression value of the gene.

### 3.3 Standardizing Network

Because the distributions of the edge weights of the three tissue-specific PPI networks were not the same, we used the following equation to standardize the weights of the three networks:

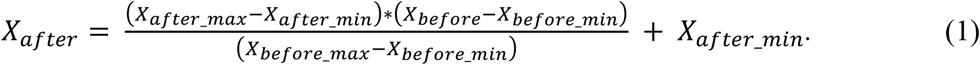

*X_after_max_* and *X_after_min_* represented the maximum and minimum of the normalized interval, respectively. Here, they were set to be 1 and 0.1 respectively. *X_before_max_* and *X_before_min_* represented the maximum and minimum of the dataset before the normalization, respectively. *X_before_* and *X_after_* represented network edge weights before and after normalization, respectively.

### 3.4 Selecting differential gene sets

For the gene expression profiles under disease conditions (leukemia, breast cancer, or prostate cancer) or under the action of drug TSA, we firstly used the z-score formula to standardize them as follows:

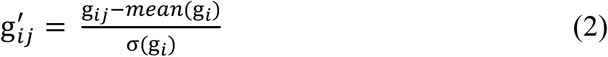

where g_*ij*_ represented the expression value of gene *i* in sample *j*, and *mean*(g_*i*_) and σ(g_*i*_) respectively represented the mean and standard deviation of the expression vector of gene *i* across all samples.

Second, we used the Limma [51] package in R to analyze the difference of standardized gene expression (see formula(2)). The log*FC* value was used to evaluate the differential expression of genes, which was defined as follows:

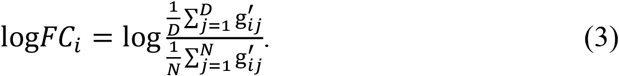

*D* and *N* represented the total number of case and control samples, respectively, and 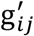 represented the expression value of gene *i* in sample *j*. If log *FC_i_* >0, gene *i* was up-regulated in case samples relative to control samples. If log*FC_i_*=0, gene *i* had no difference in two types of samples. If log*FC_i_*<0, gene *i* was down-regulated in case samples relative to control samples.

Finally, we respectively obtained three sets of differentially expressed genes under three different diseases states (leukemia, breast cancer, or prostate cancer) and a set of differentially expressed genes under the action of TSA. For a gene, if its logFC value in a disease state (leukemia, breast cancer, or prostate cancer) was negatively correlated with that under the action of TSA, the gene was selected. In this way, we obtained three gene sets acting under TSA for treating leukemia, breast cancer, and prostate cancer, which included 824, 1213, and 1160 genes respectively.

### 3.5 Mining modules from multi-layer network

In this paper, we used a tensor-based computational framework for mining recurrent heavy subgraphs (RHSs) in multilayer networks, as proposed by Li, et al. [52]. Described simply, a tensor is a multidimensional matrix, and the matrix is a second-order tensor. For any given *m* networks with the same *n* nodes that have different topologies, the following third-order tensor *A* can be used to represent it:

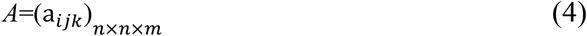

Where *a_ijk_* indicates the weight of the edge between vertices *i* and *j* in the *k^th^* network. Tensors can be used to convert discrete problems into continuous problems in graph theory. Candidate drug target modules in multi-layer network based on tensor recognition can be identified by heaviness [52]. The heaviness of a RHS is defined as the average weight of all edges in the RHS [52], which is calculated as follows:

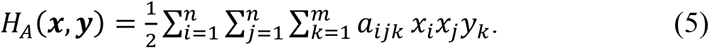

The gene vector ***x*** = (*x*_1_… *x_n_*)^*T*^, where *X_i_* = 1 if gene *i* belongs to the RHS, and *x_i_* =0 otherwise. The network vector ***y*** = (*y*_1_… *y_n_*)^*T*^, where *y_j_* =1 if the RHS appears in networkj, and *y_j_* =0 otherwise.

In this paper, we used the RHSs that have the greatest heaviness as the candidate drug-target modules of TSA.

### 3.6 Quantifying the overlap between modules

The modules containing at least three nodes were conserved. We used the following measure to quantify the overlapping coefficient *c* between two modules A and B:

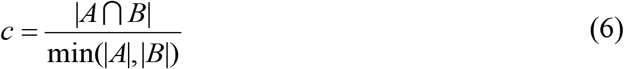

When 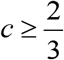, we defined the two modules A and B were overlapped modules. It will be used to measure the degree of similarity between two modules.

## 4 Results

### 4.1 Selecting three tissue-specific networks

For this work, we considered that the edges with high weights in three tissue-specific networks downloaded from the GIANT database. Moreover, to ensure the three networks had a similar density, we selected 173072 edges (top 0.5%; weights of edges approximately greater than 0.534361) from the blood network, 170017 edges (top 0.25%; weights of edges approximately greater than 0.375826) from the breast network, and 167211 edges (top 0.25%; weights of edges approximately greater than or equals to 0.31895) from the prostate network. The three tissue-specific networks (blood, breast and prostate) contained 6578, 7915 and 7844 nodes, respectively, as shown in Figure 2. They had 5484 common nodes. 434 nodes existed only in the blood PPI network, accounting for only 6% of the total number of nodes; 819 nodes existed only in the breast PPI network, accounting for only 10% of the total number of nodes; and 888 nodes existed only in the prostate PPI network, accounting for only 11% of the total number of nodes. Therefore, the three networks satisfied the principle that most genes are expressed in different tissues, and only a small portion are tissue-specific [53].

**Figure 2.**
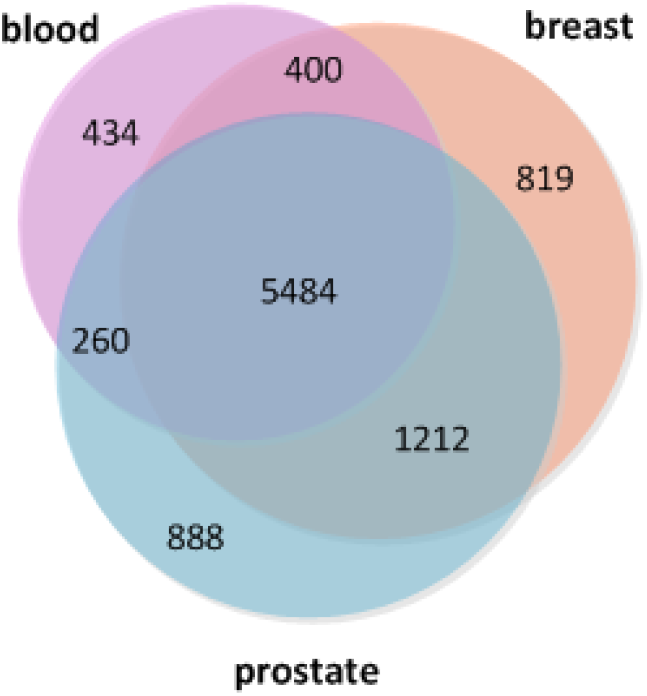
Overlapping of gene sets from three tissue-specific networks: blood, breast and prostate.

PPI networks are subject to the general distribution of scale-free networks [54], with only a few nodes having a large degree and most nodes having a relatively small degree of distribution. Here, the degree of a node in a network is the number of connections it has to other nodes. We calculated the node degree distributions of the selected three tissue-specific PPI networks, as shown in Figure 3. From Figure 3, we can see a small number of nodes have a degree of greater than 500, and these nodes are called hub nodes. Next, experiments and analyses were conducted for the filtered tissue-specific PPI networks.

**Figure 3.**
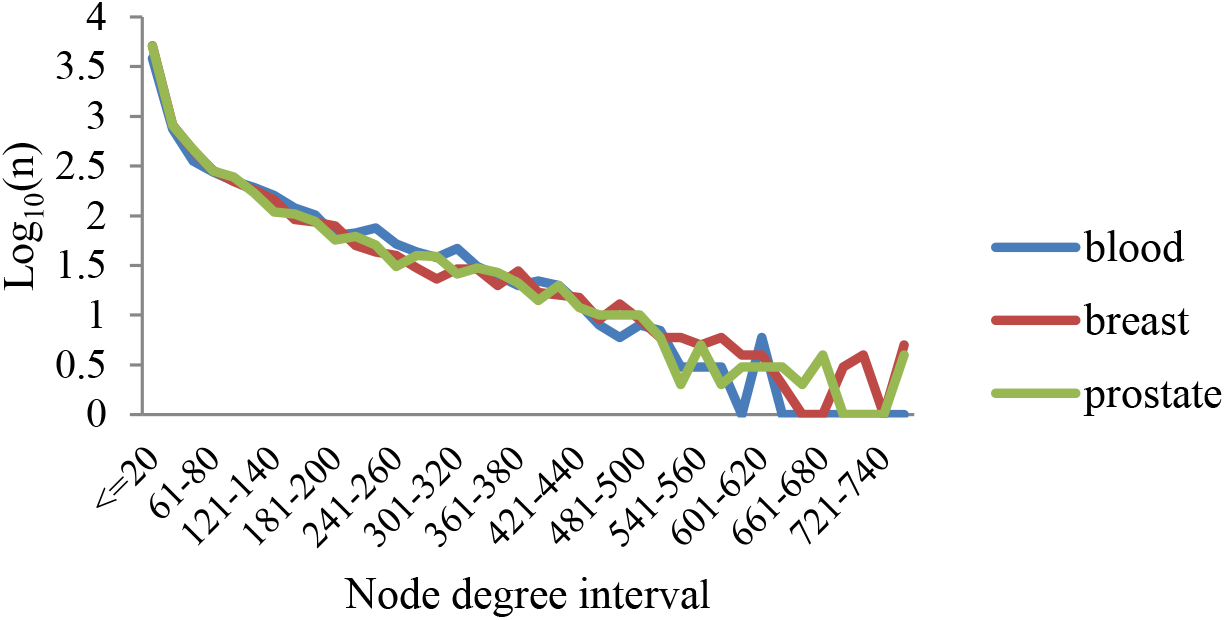
Node degree distributions of the three tissue-specific PPI networks. The X axis represents the distribution interval of the node degree. The Y axis represents Log_10_(*n*), where *n* is the number of nodes.

### 4.2 Selecting parameter *heaviness*

For the tensor-based computational framework for mining RHSs in multilayer networks, one difficulty is determining the value of the key parameter called *heaviness[52]*. We know that as the value of *heaviness* increases, the weight of the internal gene connection of the extracted candidate drug-target module becomes greater. The closer the internal connection of the module, the more reliable it is. They are more likely to perform some kind of similar function together [55]. The candidate drug-target modules extracted here are those being affected directly or indirectly by drugs.

In order to obtain the appropriate heaviness value, we gave the number of modules that can be enriched significantly in GO terms and KEGG pathways under different heaviness values, shown in Figure 4 (a) and (b) respectively. As heaviness grew, the number of enriched modules in GO and KEGG, denoted as Ne, continued to decline, but the ratio *R_ea_* between *N_e_* and the total number of modules, denoted as *N_a_*, were increasing (red curve with triangular type in Figure 4), i.e. *R_ea_* = *N_e_/N_a_*. We found when the value of *heaviness* was near 0.41, the value of *R_ea_* changed smoothly, and when *heaviness*=0.41, the number of enriched modules *N_e_* was slightly higher than its neighbor area.

**Figure 4.**
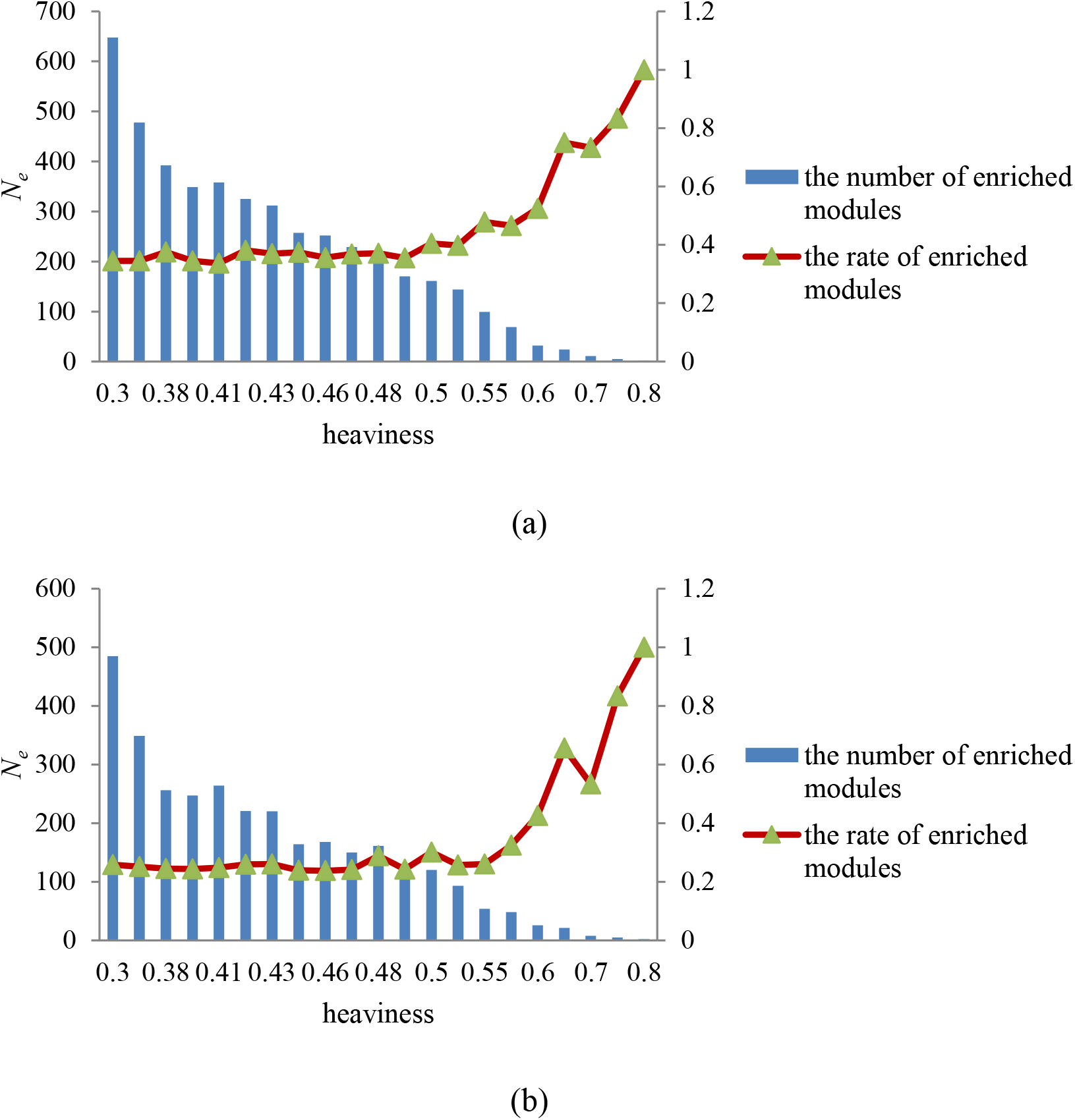
Module distribution that can be significantly enriched in the GO terms or KEGG pathways. Figure 4(a) and (b) indicated GO terms and KEGG pathways analyses, respectively. The blue bar represented the number of modules that were significantly enriched in GO terms or KEGG pathways under different values of heaviness, i.e. *N_e_*. The *Y* coordinate on the left corresponded to the change of *N_e_*. The red curve with triangular type indicated the ratio *R_ea_* between *N_e_* and the total number of modules *N_a_*. The *Y* coordinate on the right corresponded to the value of *R_ea_*.

In addition, if the more known targets of the drug appear in the module, the more likely the module is to be a potential module target. Figure 5 showed the number of overlaps between the genes included in the extracted modules and the genes that TSA affected in three different diseases. The X axis represents the value of *heaviness* (marked as “module density”), and the Y axis indicates the number of overlapping genes (marked as “number”). We found that as the value of “module density” increased, the value of “number” decreased. Compared with neighboring parameters, when the value of “module density” was 0.41, the value of the “number” for leukemia corresponding to blood tissue tended to increase (red line in Figure 5). For prostate cancer corresponding to prostate tissue (green line in Figure 5), there is a similar phenomenon.

**Figure 5.**
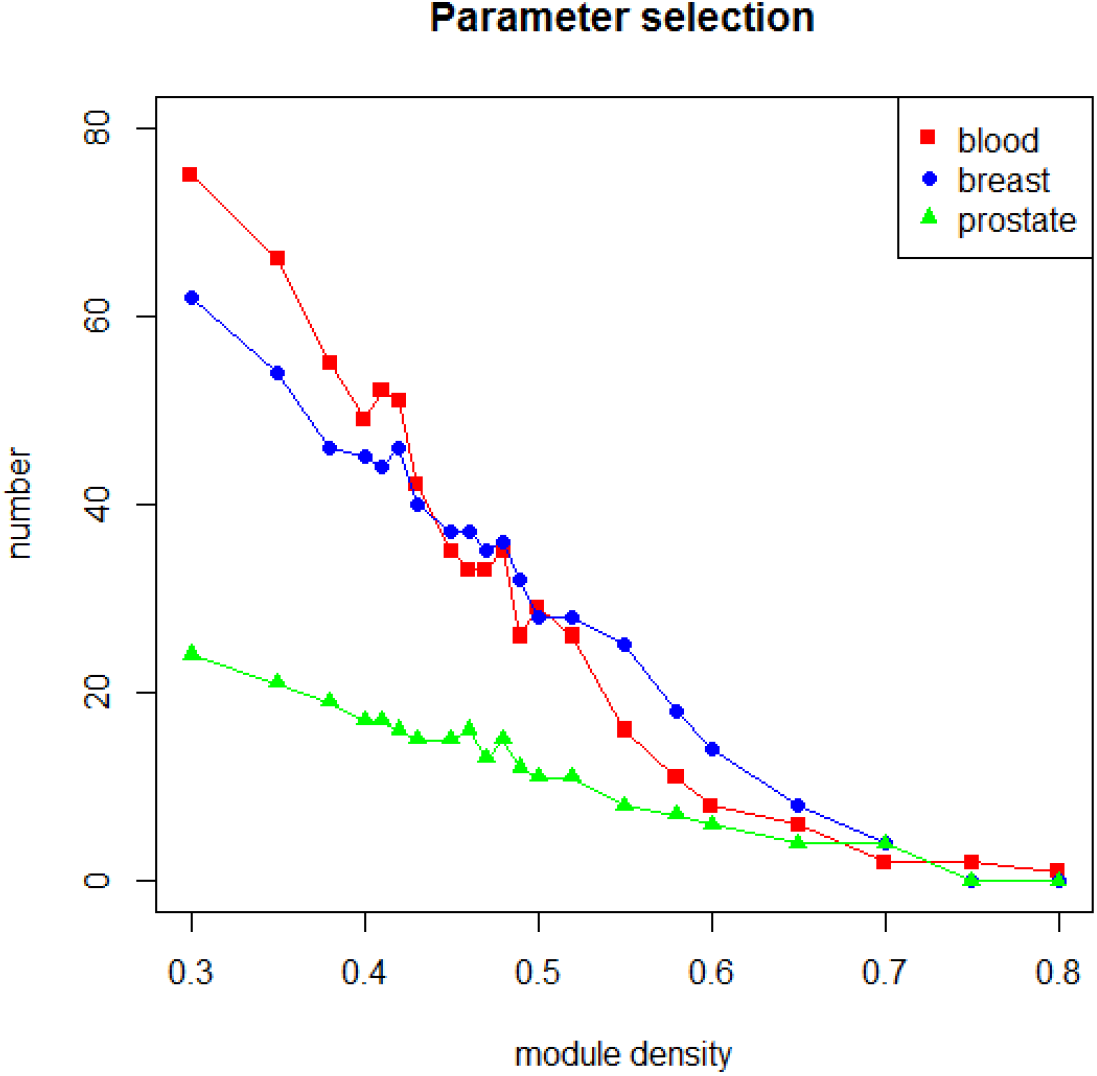
Overlapping between genes in modules generated by different values of *heaviness* and genes affected by TSA in different diseases. Red line: blood represented leukemia; Blue line: breast represented breast cancer; Green line: prostate represented prostate cancer. The X axis represented the value of *heaviness* (marked as “module density), and the Y axis indicated the number of overlapping genes (marked as “number”).

Therefore, we chose 0.41 as the value of *heaviness* and the drug-target modules obtained under this value was analyzed in the following sections.

### 4.3 Comparing predicted modules between the multi-layer and single-layer networks

#### Module overlapping comparison

In this section, based on the same module mining algorithm (see Methods section), we compared the module overlapping between the multi-layer and single-layer networks. The definition of overlap between two modules was shown in equation (6). The comparing results were shown in Figures 6 and 7. In Figure 6, when the value of the x-axis was equal to blood, breast or prostate, its corresponding y-axis value represented the number of overlapping modules between the blood, breast or prostate network and the multi-layer network. If the value of the x-axis was a combination of multiple single-layer networks, such as blood-breast or blood-breast-prostate, its corresponding y-axis value represented the number of common modules between these networks. For example, when *x*=blood-breast-prostate, its corresponding *y*-axis value was 560, which meant there were 560 common modules in blood, breast and prostate PPI networks. Figure 7 showed the proportion of the total number of modules in Figure 6 in the total number of modules mined in the multi-layer network under the same x value. And, the red square in Figure 7 indicated the proportion of 28% (corresponding to blue bar) of the overlapped modules in 560 modules. Its *y*-axis value is 0.51, which meant that 285 (560×0.51=285) common modules of the three single-layer networks can be found from the multi-layer network. Through the comparison shown in Figures 6 and 7, we found based on the multi-layer network mining functional module, not only can modules in most single-layer networks be detected, but also new modules can be discovered.

**Figure 6.**
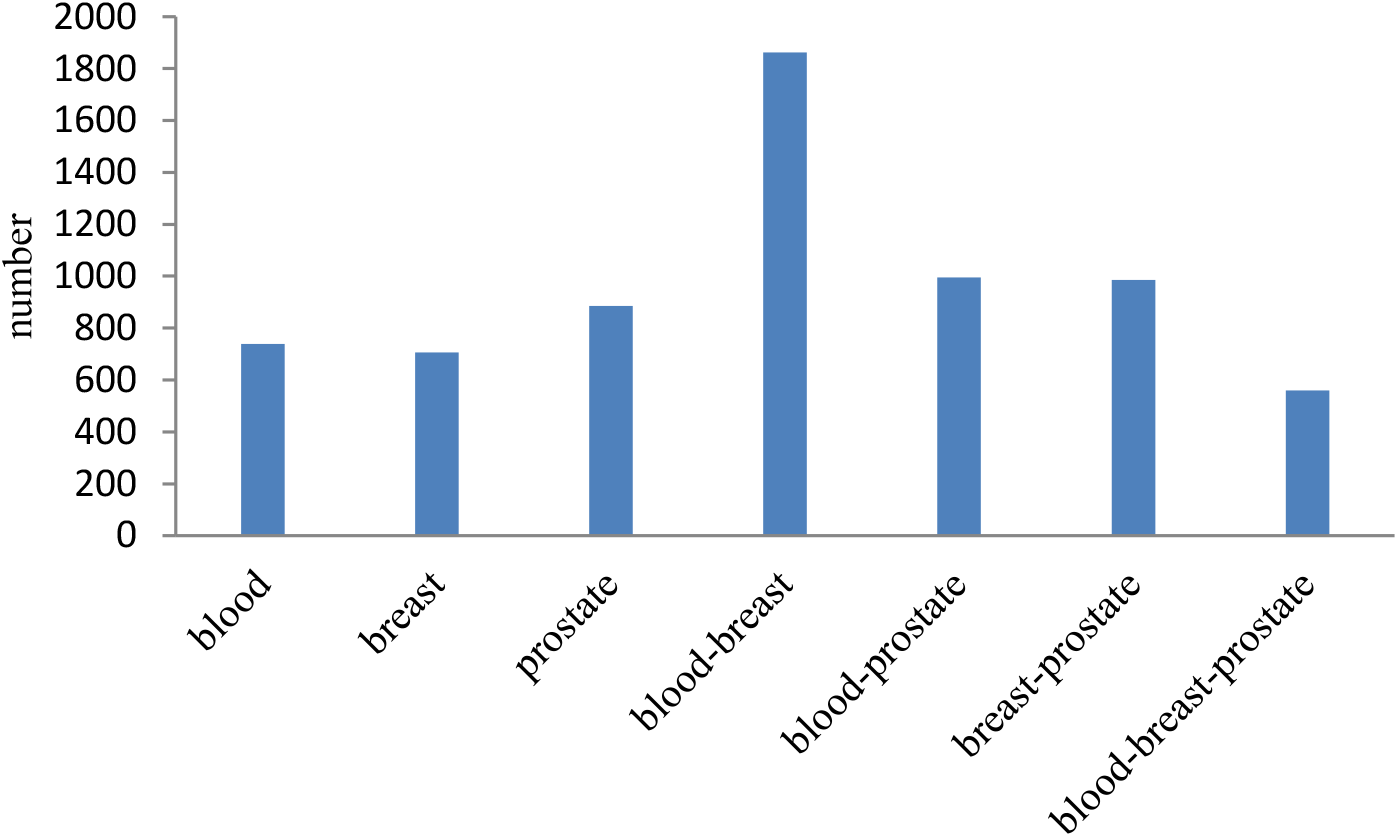
The number of overlapped modules between single-layer vs multi-layer, single-layer vs. multiple single-layer networks.

**Figure 7.**
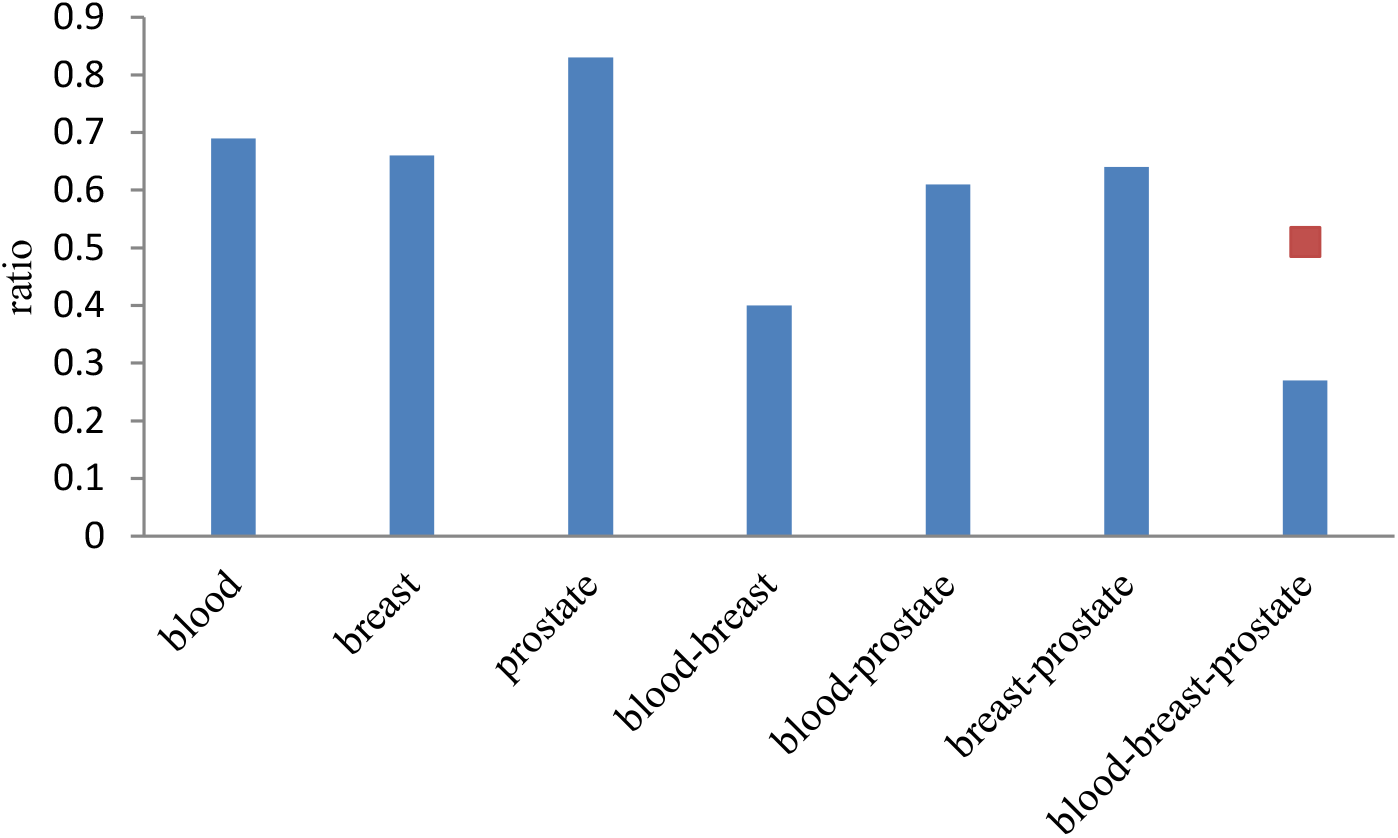
The proportion of the total number of modules in Figure 6 in the total number of modules mined in the multi-layer network under the same x value. The red square indicated the proportion of 28% (corresponding to blue bar) of the overlapped modules in 560 modules.

#### Functional enrichment comparison

We enriched the functions of modules mined in the multi-layer network and three single-layer tissue-specific networks (blood, breast and prostate) separately using GO terms [56], KEGG pathways [57], BioCarta pathways, and Reactome. Figure 8 showed a comparison of the proportions of functionally enriched modules obtained in different networks, marked as blood, breast, prostate and multi-layer. Except for BioCarta pathways, the enrichment proportions of multi-layer modules were all higher than that of single-layer modules. For BioCarta pathways, the enrichment proportion of multi-layer modules, although not as high as that of the modules in the prostate network, was slightly higher than the other two single-layer networks: blood and breast.

**Figure 8.**
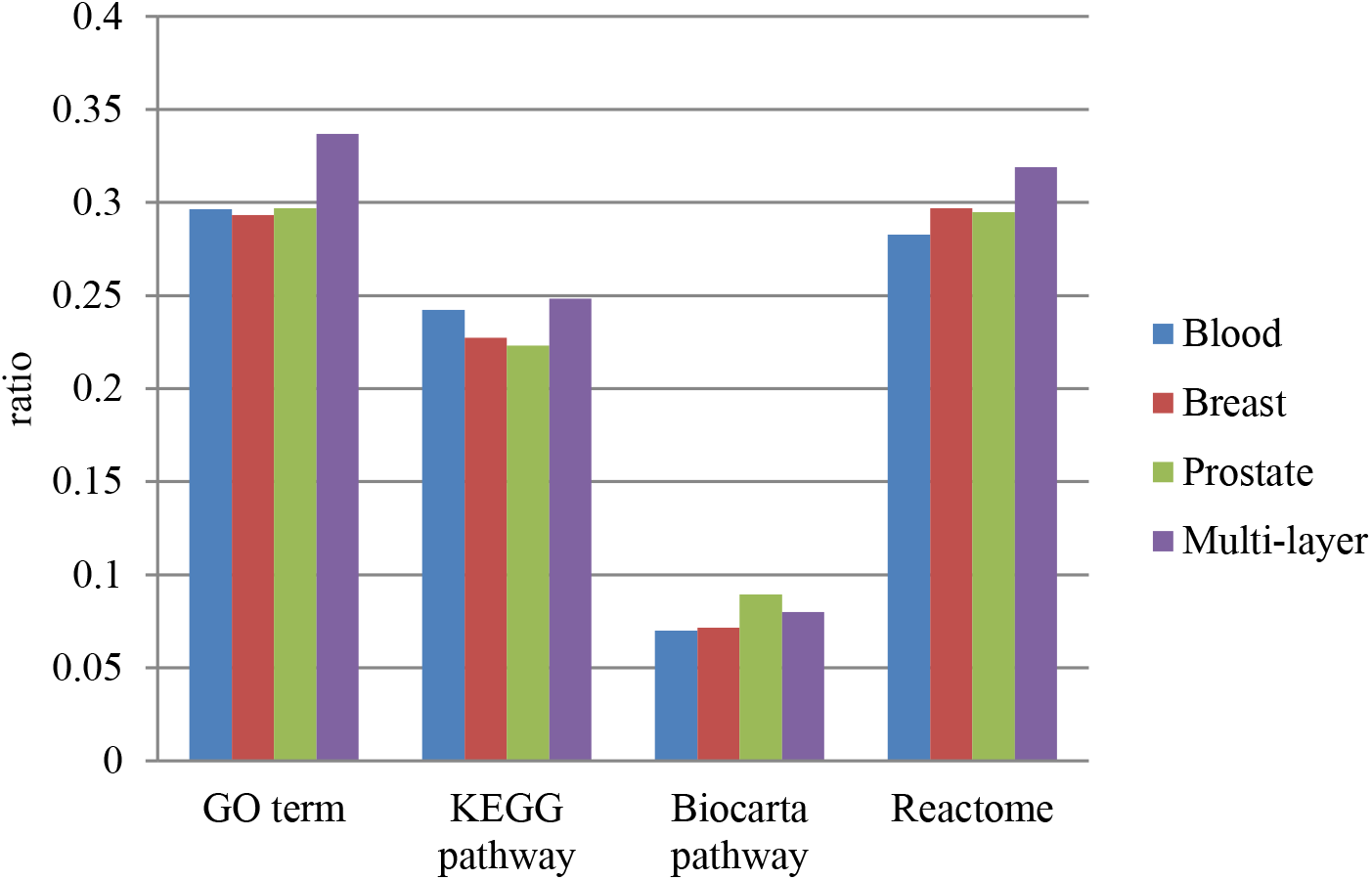
The comparison of the proportions of functionally enriched modules obtained in different networks, marked as Blood, Breast, Prostate and Multi-layer in four enrichment aspects: the GO terms, KEGG pathway, BioCarta pathway and Reactome.

We can see a higher proportion of functional modules can be predicted based on the multi-layer network than the single-layer networks.

### 4.4 Selecting modules extracted from multi-layer network

Totally, we found 1063 drug-target modules from the multi-layer network. Most of them are not meaningful. We filtered them based on the sequential three steps: the action of trichostatin A (TSA), GO enrichment and KEGG pathway.

#### 4.4.1 Based on the action of trichostatin A (TSA)

If a drug-target module contained genes that TSA can affect for leukemia, breast cancer and prostate cancer, it would be persevered for the next step analysis. Based on this principle, 26 modules were selected, shown in Table 1. For convenience, we numbered them from M1 to M26.

**Table 1.** Selected candidate drug-target modules based on genes affected by TSA.

| Module ID | Entrez ID of genes in modules |
| --- | --- |
| M1 | 890 7153 4085 6241 701 22974 6790 3161 11130 10403 6240 10051 51203 1434 1719 3832 7298 5984 10592 4173 891 9319 2237 3838 990 47 90 87 |
| M2 | 7520 142 1019 5111 5591 6749 2237 5036 4522 6241 4175 10606 5982 1736 |
| M3 | 22948 10213 10969 471 1434 3329 5686 1503 9221 908 5901 5036 3838 7371 |
| M4 | 5901 7334 7520 7443 10576 7153 10213 26135 6636 6427 5902 6428 6240 |
| M5 | 22948 7203 6950 10574 11222 1164 4830 7334 |
| M6 | 4172 6627 1503 10528 11130 2237 7398 9521 5985 |
| M7 | 6426 4436 10772 10236 3838 26135 1665 23165 10576 7520 |
| M8 | 7153 5557 6790 672 8317 10733 4001 1736 |
| M9 | 6426 9221 6434 7334 3015 1736 2237 3184 2956 6427 |
| M10 | 10574 158 7965 142 1503 7411 4176 1736 8607 7203 5901 5902 |
| M11 | 6637 5111 3148 3182 6434 |
| M12 | 1434 3308 908 4869 6950 7203 3336 3838 |
| M13 | 10492 1503 3182 |
| M14 | 3276 5725 3609 6597 4176 6627 |
| M15 | 6194 6124 6201 6137 11224 6143 6193 6217 6152 6139 6136 6161 23521 6133<br>6175 4736 6207 6218 6135 6128 6146 3646 1933 47 87 39 29 90 95 |
| M16 | 3014 84823 6597 5036 |
| M17 | 3065 142 1786 6597 |
| M18 | 3066 3065 5928 2146 6597 |
| M19 | 86 6597 10856 |
| M20 | 6597 6599 5591 4173 4172 |
| M21 | 6597 23246 8662 |
| M22 | 5036 10574 3182 |
| M23 | 3329 7203 6428 |
| M24 | 10606 6950 4691 3183 6741 3843 5901 |
| M25 | 890 7371 3251 1665 |
| M26 | 5557 990 9493 9833 1060 |

#### 4.4.2 Based on GO enrichment

In this section, the 26 selected modules will be processed further. Gene Ontology [56] (GO) is a framework for biological models that divides genes based on three different aspects: molecular function (MF), cellular component (CC) and biological processes (BP). A module will be chosen if the module has overlapping GO terms with the drug-target gene. Further, we also needed to make sure that the module was related to the disease, so the module should have overlapping GO terms with disease-causing genes. Therefore, we validated the 26 selected modules using the following steps.

##### Step 1: Filtering modules by overlapping GO terms with drug targets

We downloaded the targets of TSA through SuperTarget (http://insilico.charite.de/supertarget/), a database developed to collect information about drug-target relations [58], and DrugBank (https://www.drugbank.ca/), a unique bioinformatics and cheminformatics resource that combines detailed drug data with comprehensive drug target information databases [59]. TSA targets were mapped to DAVID (version 6.8) [60, 61] for GO enrichment analysis. TSA targets enriched in 74 GO terms, including 42 BP terms, 20 MF terms and 12 CC terms. For each module in the selected 26 modules, the same analysis in DAVID will be made to them. That is to say, each module corresponded to a GO term list, including BP, MF and CC terms.

For each of BP, MF and CC, the ratio of the number of overlapping terms between each module and TSA targets to the number of terms enriched by the module was calculated. The results were shown in Figure 9.

**Figure 9.**
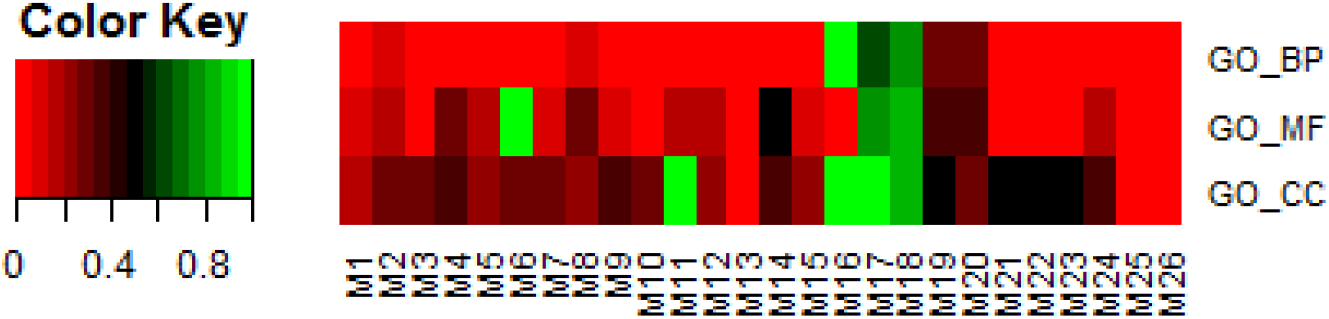
The ratio of the number of overlapping terms between each of the 26 selected modules and TSA targets to the number of terms enriched by the module in BP, MF and CC respectively.

If a module had more overlapping GO terms in all the three aspects (BP, MF and CC), it will be conserved in this step for further analysis. Eight modules were selected from the 26 modules in this step: M1, M2, M3, M8, M17, M18, M19 and M20.

##### Step 2: Filtering modules by overlapping GO terms with disease genes

In order to improve the reliability of the drug module target, we need to further filter the eight candidate modules got from step 1 based on disease genes. The genes related to leukemia, breast cancer and prostate cancer were got from OMIM [62, 63] and GWAS database [64]. We projected these disease genes into DAVID for GO enrichment analysis. For three diseases: leukemia, breast cancer and prostate cancer, the overlap rate of GO terms in BP, MF and CC enriched by disease genes and the selected eight modules (M1, M2, M3, M8, M17, M18, M19 and M20) were shown in Figure 10(A) to (C) respectively.

**Figure 10.**
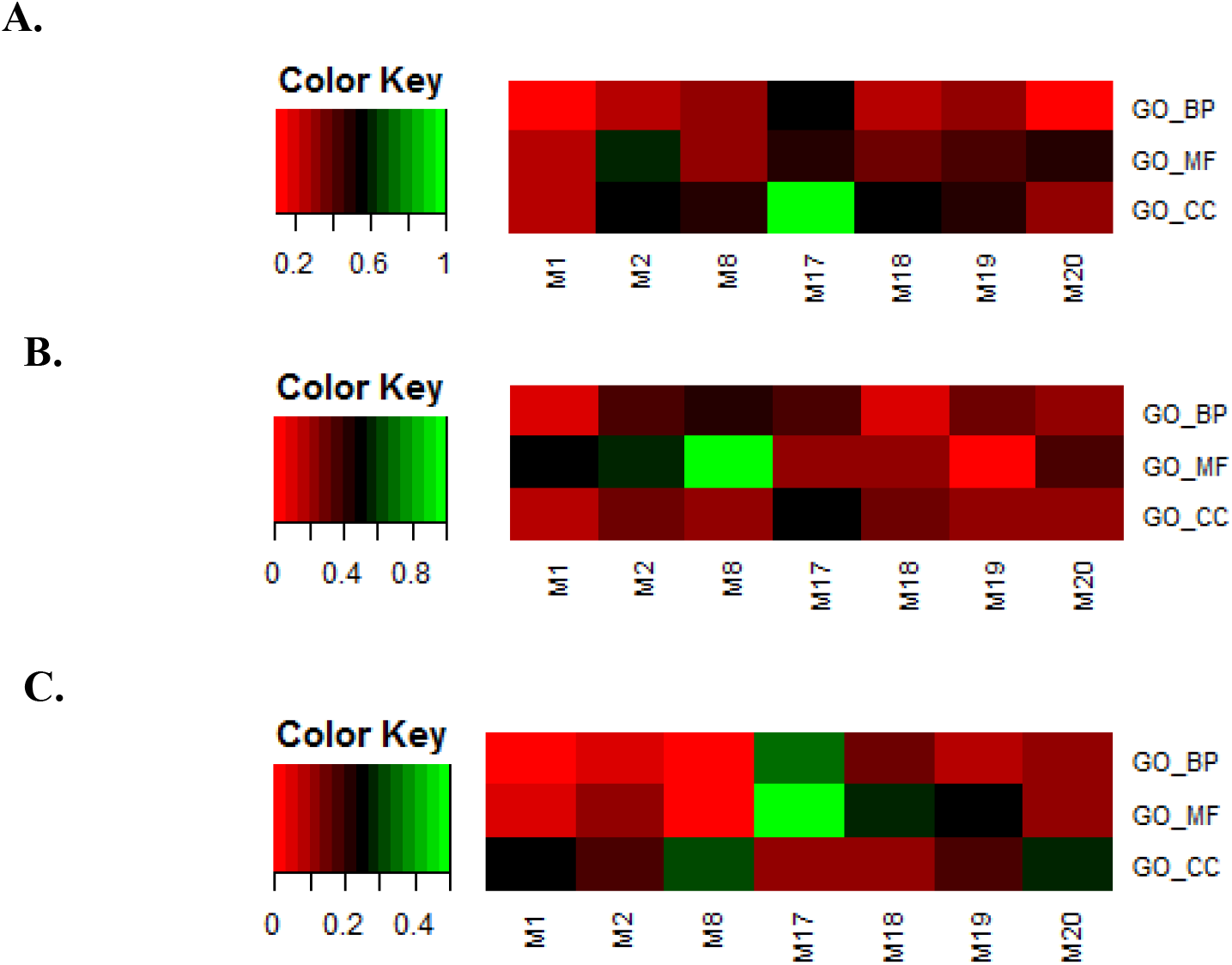
The overlap rate of GO terms in BP, MF and CC enriched by disease genes and the selected eight modules in three diseases. (A) leukemia (B) breast cancer (C) prostate cancer

Only those modules were conserved, which overlapped with all the three disease gene sets in each of the three aspects: BP, MF and CC. In this way, we obtained four modules: M2, M17, M18 and M20.

#### 4.4.3 Based on KEGG pathways

To further indicate the relationship between modules, drugs and diseases, we also analyzed the connections between internal genes of modules and their first-order neighbors outside the modules. The tighter the gene is connected to the module, the more it is affected by the module. The strength of the connection between a first-order gene *j* and a module *D* was defined as a score *score*(*j,D*) in formula (7):

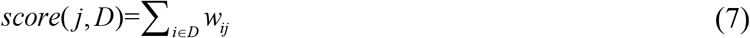

Here, *w_ij_* represented the edge weight between *j* and *i* (*i* ∈ *D*).

For each of the four modules: M2, M17, M18 and M20, we separately calculated their connection strength with first-order nodes in blood, breast and prostate tissue-specific PPI networks. The results were shown in Figure 11 A to C. The *X* axis represented the score value of connection strength. The *Y* axis indicated the number of first-order genes of modules. For each module, we selected its first-order genes with high scores to analyze the KEGG pathway enrichment by DAVID tool [57, 65].

**Figure 11.**
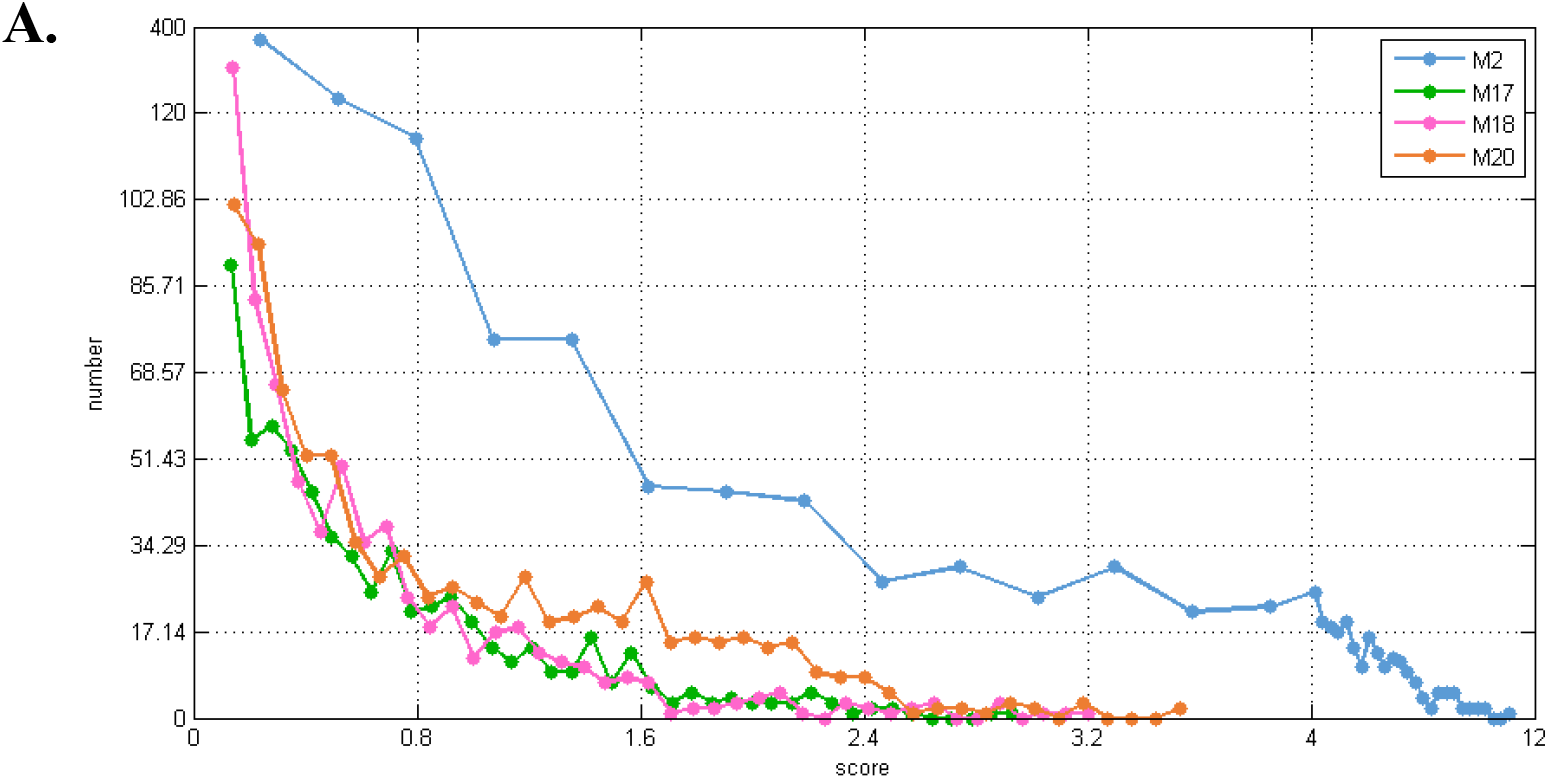
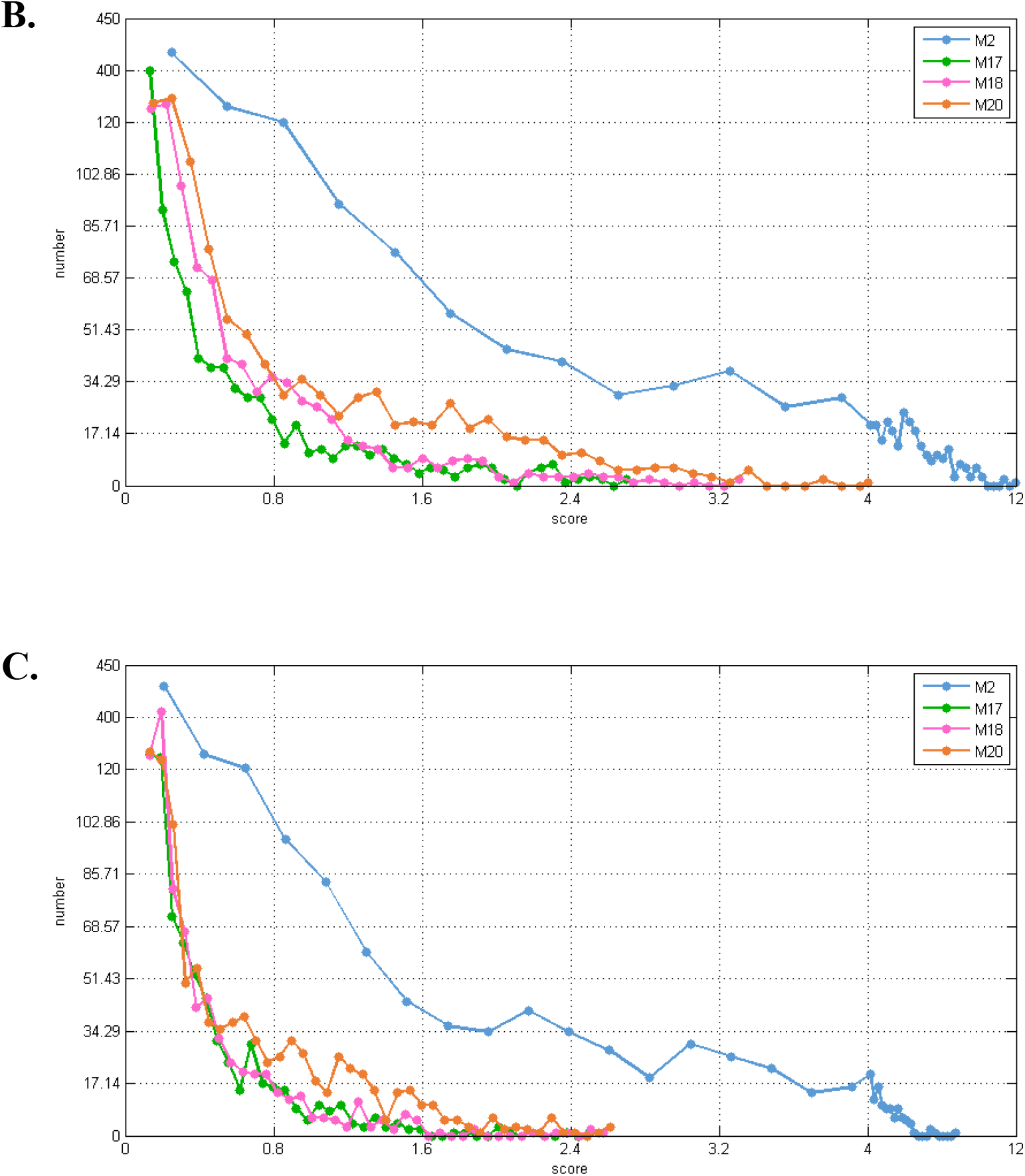
Distribution of connection strength between four modules and their corresponding first-order genes in three tissue-specific networks. The *X* axis represented the score value of connection strength. The *Y* axis indicated the number of first-order genes of modules. A, B, and C indicated the blood, breast, and prostate tissue-specific PPI networks, respectively.

In different networks, the distribution of first-order neighbor nodes of the same module was similar. Different numbers of nodes in each module caused them to have different numbers of first-order neighbors. That is to say, the four modules have different first-order neighbors in the same network. We only kept first-order neighbors with high scores for KEGG Pathway enrichment analysis. We used the following method to filter the first-order neighbors for each module. First, we selected 1,000,000 genes randomly in the first-order genes of a module and got their connection strength scores. We sorted all the scores in descending order and took the 50th value as the threshold. Finally, those genes whose scores were greater than this threshold were conserved. We also used DAVID to make KEGG pathway enrichment to these first-order neighbors. The results for blood, breast and prostate tissue-specific networks were shown in Table 2 to 4 respectively. For example, in Table 2, for module M2, we got the threshold was 7.0 and 58 first-order genes were conserved. The 58 genes were enriched to KEGG pathways database by using DAVID and 12 significant pathways were obtained.

**Table 2.**
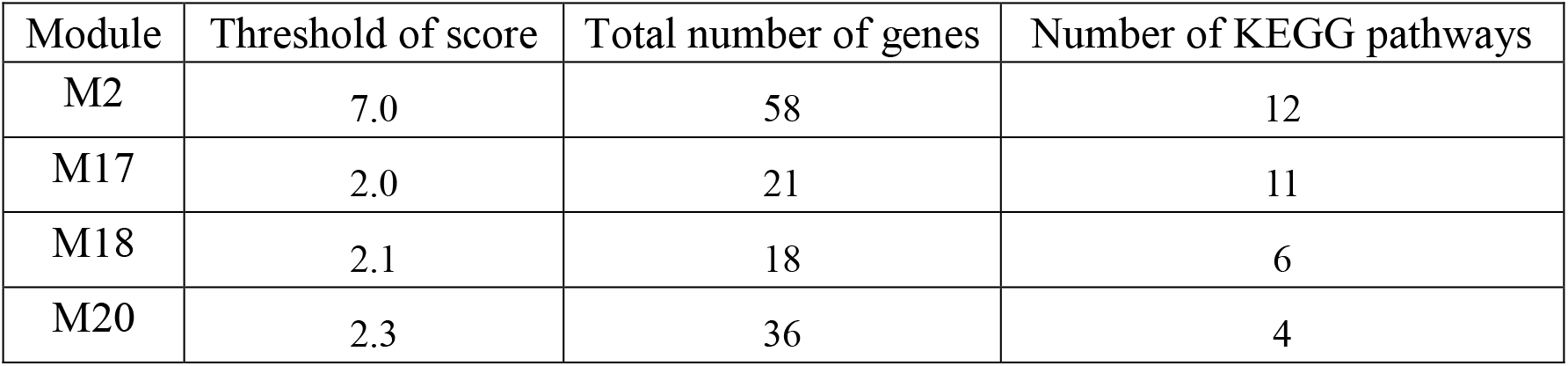
KEGG pathway enrichment of conserved first-order genes of modules in blood tissue-specific PPI network.

**Table 3.**
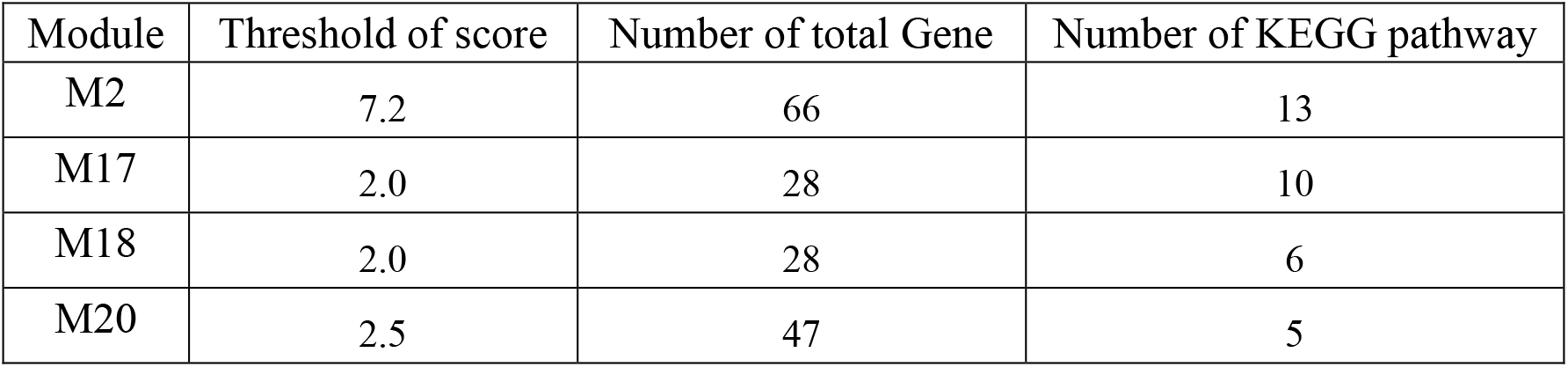
KEGG pathway enrichment of conserved first-order genes of modules in breast tissue-specific PPI network.

**Table 4.**
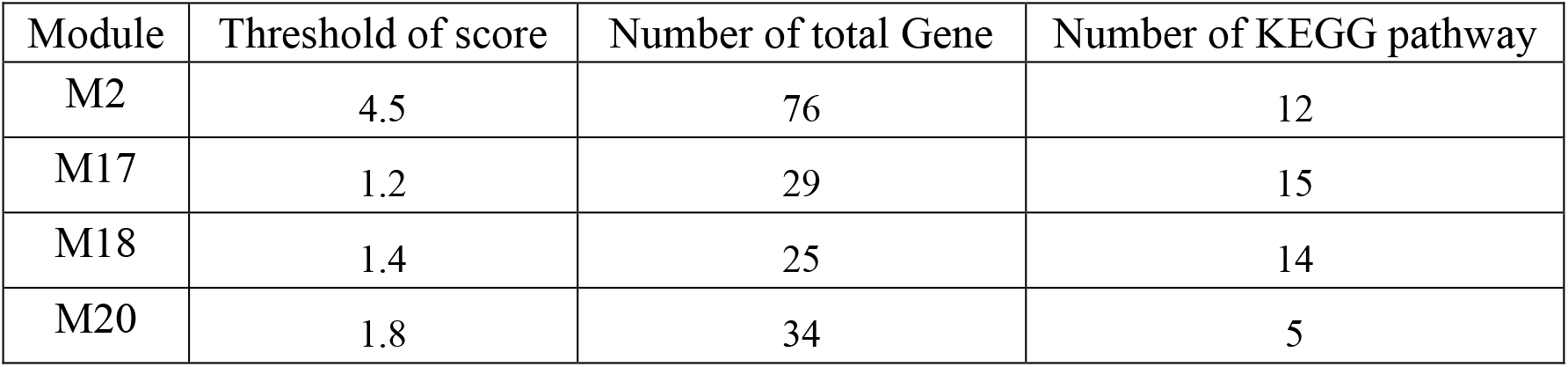
KEGG pathway enrichment of conserved first-order genes of modules in prostate tissue-specific PPI network.

| Module | Threshold of score | Number of total Gene | Number of KEGG pathway |
| --- | --- | --- | --- |
| M2 | 4.5 | 76 | 12 |
| M17 | 1.2 | 29 | 15 |
| M18 | 1.4 | 25 | 14 |
| M20 | 1.8 | 34 | 5 |

Drugs can affect the module’s first-order neighbors by affecting the module for disease treatment. The higher the overlap of KEGG pathways between first-order genes of modules and targets of drug TSA, the better the treatment effect. Table 5 showed the number of overlapping KEGG pathways between first-order genes of modules and targets of drug TSA. Finally, two modules (M17 and M18) were conserved.

**Table 5.** The number of overlapping KEGG pathways enriched by first-order genes of modules and targets of drug TSA in three tissue-specific PPI networks.

| Tissues | M2 | M17 | M18 | M20 |
| --- | --- | --- | --- | --- |
| Blood | 0 | **3** | **1** | 0 |
| Breast | 1 | **4** | **1** | 0 |
| Prostate | 1 | **4** | **4** | 1 |

We also analyzed the overlapping KEGG pathways between the first-order genes of modules M17 and M18 and causing genes of leukemia, breast cancer and prostate cancer. We found they had more overlapping. Table 6 gave the number of overlapping KEGG pathways. The results, on the other hand, verified that TSA can treat these three diseases and the two module were feasible.

**Table 6.** The number of overlapping of KEGG pathways enriched by first-order genes of modules and causing genes of three diseases.

| Cancer | M17 | M18 |
| --- | --- | --- |
| Leukemia | **6** | **1** |
| Breast cancer | **3** | **1** |
| Prostate cancer | **1** | **1** |

For the overlapping KEGG pathways enriched by these two modules and drug TSA, we found that they had the highest proportion of chronic myeloid leukemia pathways, which were closely related pathways for leukemia. This showed indirectly that TSA can act on the disease by acting on the M17 and M18 modules, thereby proving the effectiveness of using our method STAM for finding drug-target modules. That is to say, M17 and M18 were the potential treatment patterns of TSA.

### 4.5 Validating and analyzing the significance of M17 and M18

The tissue-specific PPI networks were obtained from GIANT. The weights on the edges represented confidence: the greater the weight, the higher the confidence and the closer the interactions between genes. To ensure that the drug-target modules we found, i.e. M17 and M18, were not random, the following analysis from step 1 to step 4 was performed on modules M17 and M18 separately in blood, breast, and prostate tissue-specific PPI networks.

Step 1: The sum of the weights of the inner edges of module *m* in a tissue-specific PPI network was calculated and saved as *S_m_*.

Step 2: The tissue-specific PPI network used in step 1 was randomly disturbed. The network connections were unchanged, and the weights of the edges were disturbed randomly.

Step 3: In the randomized network obtained in step 2, the sum of the weights of the inner edges of module *m* was calculated again.

Step 4: Steps 2 and 3 were repeated 10^6^ times, and the results were saved as 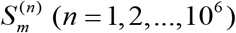.

Step 5: The results 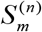 obtained in step 4 were compared with the result *S_m_* obtained in step 1. The formula was as follows:

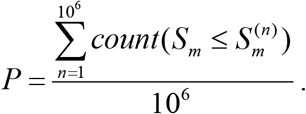

where *P* denoted the p-value of the drug-target module *m*. If 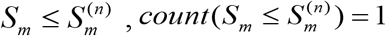; otherwise, 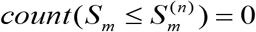. The smaller the *p*-value, the more meaningful of the module *m*. Based on the above analysis process (step 1 to step 5), M17 and M18 were respectively analyzed in three tissue-specific PPI networks to obtain their corresponding *p* values, and the results were shown in Table 7.

**Table 7.** Significance of target modules M17 and M18 of TSA in three tissue-specific PPI networks.

| Tissues | p-values of M17 | p-values of M18 |
| --- | --- | --- |
| Blood | 6.27e-04 | 6.36e-06 |
| Breast | 3.24e-04 | 0 |
| Prostate | 1.64e-04 | 0 |

The *p*-values of M17 and M18 in blood, breast, and prostate tissue-specific PPI networks were all less than 0.05. The results demonstrated that the target modules M17 and M18 of TSA were highly statistically significant, and their internal genes had strong interactions.

To further verify that M17 and M18 were also highly significant in other TSA-related tissue-specific PPI networks, we obtained the other five cancers treated by TSA, marked as “T”, from the Comparative Toxicogenomics Database (CTD)[66]. The five cancers were lung cancer, colon cancer, ovarian cancer, pancreatic cancer and myeloma. Their related tissue-specific networks were downloaded from GIANT, and edges with low weights were deleted. The process was consistent with the tissue-specific PPI network preprocessing of blood, breast and prostate cancer tissue in section 4.1. In the five processed networks, we calculated the p-values of module targets M17 and M18. The results were shown in Table 8.

**Table 8.** Significance of target modules of TSA in other five TSA-related PPI networks.

| Tissues | Number of edges | The minimal edge weight | p-values of M17 | p-values of M18 |
| --- | --- | --- | --- | --- |
| Lung | 149,495 | 0.374935 | 2.12e-04 | 0 |
| Colon | 163,180 | 0.317351 | 6.91e-04 | 0 |
| Ovarian | 161,487 | 0.37902 | 2.67e-04 | 0 |
| Pancreas | 161,147 | 0.312249 | 6.58e-04 | 8.43e-05 |
| Marrow | 154,621 | 0.391356 | 0.0242 | 9.23e-04 |

From Table 8, we can see that the *p*-values of M17 and M18 in the five tissue-specific PPI networks were all less than 0.05. This findings suggested that the significance of M17 and M18 in these tissue-specific PPI networks was high, which indicated that drug TSA was likely to treat other diseases by acting on M17 and M18. Therefore, there was strong evidence to support that M17 and M18 were the treatment patters of TSA.

### 4.6 Analyzing differences in co-expression networks

Genetic variant-induced perturbations in network properties give rise to altered phenotypes, such as diseases [67]. In this section, we studied the different connections of M17 and M18 in co-expression networks in disease and normal states. We used Pearson correlation coefficients to build co-expression network connections of genes in M17 and M18 in disease and control samples, respectively. We found that they were very different, shown in Figures 12 and 13. Figures 12 and 13 showed M17’s and M18’s co-expression network connection differences in normal and tumor-based states under three different conditions, respectively. For example, in the control (normal) and case (tumor) samples of leukemia (Figures 12 (A1) and (A2)), none of the gene pairs in M17 shared similar relationships in the two co-expression networks. For module M18, the connection differences between genes in case and control samples were very big, e.g., in leukemia (Figures 13 (A1) and (A2)). These results showed that the three diseases (leukemia, breast cancer and prostate cancer) were closely related to modules M17 and M18, and it was possible that the modules caused different diseases because of connection changings under different conditions. On the other hand, it also indicated that the modules M17 and M18 were biologically significant and were likely potential drug treatment patterns.

**Figure 12.**
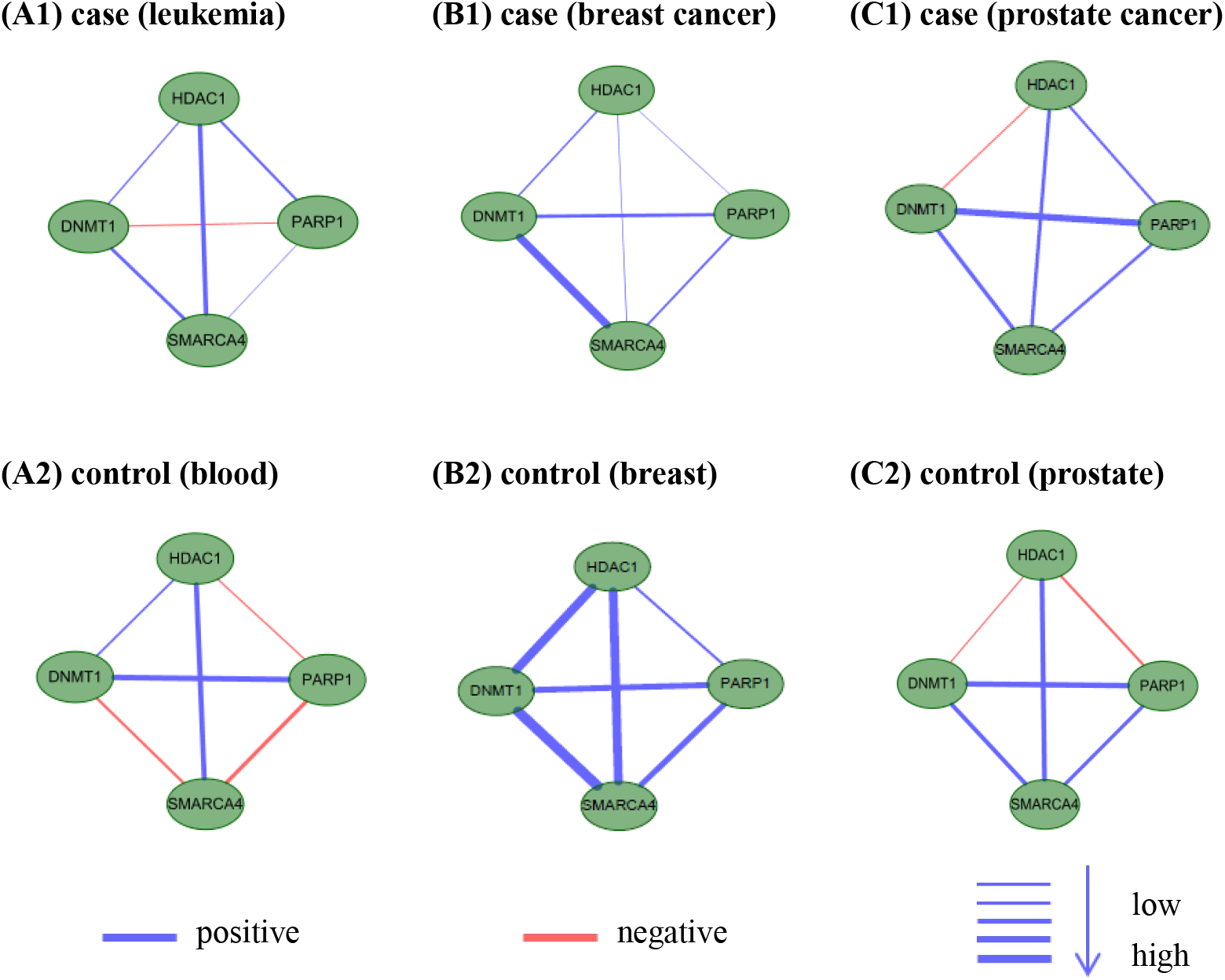
M17’s co-expression network connection differences in normal and tumor-based states under three different conditions (leukemia, breast cancer and prostate cancer).

**Figure 13.**
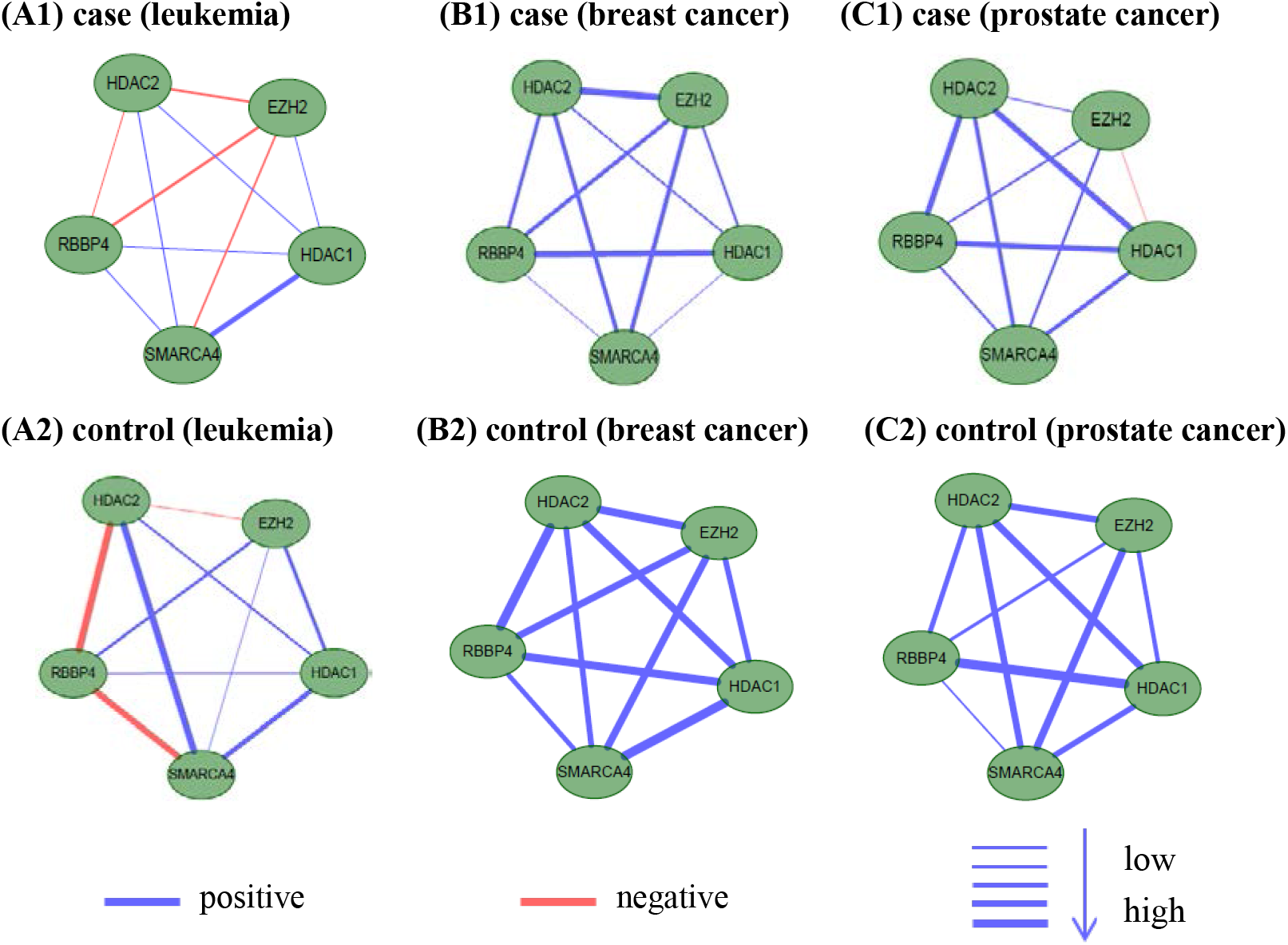
M18’s co-expression network connection differences in normal and tumor-based states under three different conditions (leukemia, breast cancer and prostate cancer).

### 4.7 PubMed literature validation for module genes

M17 contained four genes: HDAC1, PARP1, DNMT1 and SMARCA4, and M18 module contained five genes: HDAC2, HDAC1, RBBP4, EZH2 and SMARCA4. Among them, HDAC1 and HDAC2 are the targets of Trichostatin A (TSA), which is a potent inhibitor of histone deacetylase (HDAC) [33, 68]. In M17 and M18, HDAC1 and HDAC2 linked to other genes to perform different functions. For example, DNMT1 in module M17 can be translated into a homodimer and form a stable complex with HDAC1, inhibiting transcription of the E2F-responsive promoter [69]. The combination of HDAC2, HDAC1, and RBBP4 in module M18 can form part of the core histone deacetylase (HDAC) complex, while TSA chelates zinc ions in the recesses of the active site through the hydroxamic acid group of HDAC. In turn, this interaction prevents the catalytic action of HDAC [70]. The transcriptional translation of SMARCA4 in modules M17 and M18 is followed by part of the CREST-BRG1 complex, and the activity-dependent induction of NR2B expression involves the release of the HDAC1 gene. However, the CREST-BRG1 complex binds to the NR2B promoter and participates in the transcriptional activation and selection of the gene’s inhibitory process, thereby affecting the release of the HDAC1 gene [71].

To further analyze the relationship between genes in modules (M17 and M18) and TSA, we performed literature verification. The researchers found SMARCA4 (also known as BRG1) are involved in a variety of developmental processes, transcriptional regulation, DNA repair, cell cycle regulation, and cancer [72]. SMARCA4 is an ATPase subunit essential for the SWI/SNF chromatin remodeling complex in mammals, and it is also an antibody to the tumor suppressor gene SNF2β [73]. It can destroy the target region’s nucleosomes by using the energy generated by ATPase hydrolysis [73]. Mackmull et al. [74] found that SMARCA4 is affected at the protein level after 12 hours of TSA treatment, and its richness at the protein level is regulated by a combination of transcriptional and post-transcriptional mechanisms. Using gene silencing techniques, they demonstrated that a decrease in SMARCA4 richness is sufficient to regulate transcriptional changes induced by TSA[74].

PARP1 modifies various nuclear proteins by encoding a chromatin-associated poly ADP-ribosyltransferase, which is dependent on DNA and participates in a variety of important cellular processes that regulate molecular activity associated with cellular recovery from DNA damage [75]. Non-homologous end joining (NHEJ) is one of the major ways to repair DNA double-strand breaks (DSBs). TSA offers a significant increase in the probability of PARP1 binding to chromatin DSBs, and poly ADP-ribose co-localized with DSBs in TSA-bearing leukemia patients also increases significantly [62]. In addition, the knock-out of PARP1 inhibits the effect of TSA on NHEJ. These results all indicated that by taking TSA, leukemia patients increased the probability of PARP1 binding to chromatin DSBs and poly ADP ribose, thereby reducing the cytotoxicity of NHEJ and leukemia cells [76].

## 5 Conclusion

In biological networks, a drug’s therapeutic action manifested itself as a disturbance to the network. Usually, a drug can have a therapeutic effect on many diseases. That is to say, there is a pattern in the treatment of drugs, such as modules. Drugs treated different diseases by acting on a gene set with a similar network structure. Because diseases were tissue-specific, there were overlapping and different connections between same gene sets in protein-protein interaction networks associated with different diseases. Based on the above information, we used TSA as an example to establish a three-layer tissue-specific network of the three diseases it treats, and used the tensor-based multi-layer network mining algorithm [52] to identify the TSA treatment patterns. In this paper, we also named the patterns as target modules. Using the multi-layer network mining algorithm, it is possible to extract the structure of the same gene set with different connections in different layers. Finally, two potential target modules of TSA, named as M17 and M18, were found.

In this paper, we tried to find treatment patterns, i.e. target modules in multi-layer networks, for drugs through a new framework. When selecting the first-order neighbors of modules, we selected the direct neighbors of the module with high relevance scores. These genes had a greater influence on the modules, making our results more reliable. We verified our results from multiple perspectives, including difference and functional enrichment comparison, co-expression network analysis and literature verification, et al.

There are some limitations of our method. For example, the tissue-specific protein interaction networks we used were incomplete and had false positive connections. Although we have done a lot of analysis to our results, we lack biological experimental validation. In future work, we will correct these shortcomings and further improve the framework to make the results more reliable.

## 6 Acknowledgments

This work was supported by the National Natural Science Foundation of China (Nos. 61672406) and the Fundamental Research Funds for the Central Universities under Grant no. JB180307.

